# The diversity, structure and function of heritable adaptive immunity sequences in the *Aedes aegypti* genome

**DOI:** 10.1101/127498

**Authors:** Zachary J. Whitfield, Patrick T. Dolan, Mark Kunitomi, Michel Tassetto, Matthew G. Seetin, Steve Oh, Cheryl Heiner, Ellen Paxinos, Raul Andino

## Abstract

The *Aedes aegypti* mosquito is a major vector for arboviruses including dengue, chikungunya and Zika virus. Combating the spread of these viruses requires a more complete understanding of the mosquito immune system. Recent studies have implicated genomic endogenous viral elements (EVEs) derived from non-retroviral RNA viruses in insect immunity. Because these elements are inserted into repetitive regions of the mosquito genome, their large-scale structure and organization with respect to other genomic elements has been difficult to resolve with short-read sequencing. To better define the origin, diversity and biological role of EVEs, we employed single-molecule, real-time sequencing technology to generate a high quality, long-read assembly of the *Ae*. *aegypti*-derived Aag2 cell line genome. We leverage the quality and contiguity of this assembly to characterize the diversity and genomic context of EVEs in the genome of this important model system. We find that EVEs in the Aag2 genome are acquired through recombination by LTR retrotransposons, and organize into larger loci (>50kbp) characterized by high LTR density. These EVE containing loci are associated with increased transcription factor binding sight density and increased production of anti-genomic piRNAs. We also detected piRNA processing corresponding to on-going viral infection. This global view of EVEs and piRNA responses demonstrates the ubiquity and diversity of these heritable elements that define small-RNA mediated antiviral immunity in mosquitoes.

## INTRODUCTION

Arboviruses such as Dengue virus (DENV), Chikungunya virus (CHIKV), and the newly emerging Zika virus, cause widespread and debilitating disease across the globe (Bhatt et al., 2013). The primary vector of these viruses, *Aedes aegypti,* has a global tropical/subtropical distribution (Kraemer et al., 2015) creating distinct, geographically isolated populations. The genetic diversity of *Aedes* populations has resulted in differential competence for vectoring virus (Bennett et al., 2002). Differences in the insect immune system are critical factors in determining competence (Kramer, 2016; Kramer & Ciota, 2015) and recent studies suggest that virus-derived sequences in the mosquito genome may contribute to resistance (Kunitomi et al., submitted). Comparative genomics may explain these differences in vector competence between *Ae. aegypti* populations, however the repetitive nature of the mosquito genome has been refractory to assembly.

The genomic acquisition of viral sequences represents an important source of genomic diversity and immune innovation in eukaryotes (Aswad & Katzourakis, 2012; Chuong, Elde, & Feschotte, 2016; Feschotte & Gilbert, 2012; Katzourakis & Gifford, 2010). Most virus-derived sequences are retroviral and are acquired through the process of proviral genomic integration. Many examples of acquired retroviral genes evolving new functions within the host have been described (Chuong, et al., 2016). In addition, many sequences originating from non-retroviruses have integrated into the genomes of their eukaryotic hosts. These non-retroviral endogenous viral elements (EVEs) are thought to be acquired through the action of endogenous retrotransposon-derived reverse transcriptases (Belyi, Levine, & Skalka, 2010a, 2010b; Gilbert & Feschotte, 2010; Horie et al., 2010; Katzourakis & Gifford, 2010; Taylor, Leach, & Bruenn, 2010). Consistent with this model, DNA derived from RNA viruses is produced in persistently infected *Drosophila* cell lines (Goic et al., 2013) and in infected *Aedes albopictus* mosquitoes (and multiple mosquito-derived cell culture lines). Synthesis of this viral DNA (vDNA) depends on the activity of endogenous reverse transcriptases (Goic et al., 2016; Goic, et al., 2013). Furthermore, sequencing of viral DNA isolated from *Drosophila* cell lines (Goic, et al., 2013) has demonstrated the formation of recombinants between viral DNA sequences and transposable elements.

Insects rely on RNAi-based immune defenses, wherein viral dsRNA intermediates are recognized and processed through a dicer- and argonaute-mediated pathway, ultimately leading to cleavage of viral RNA and protection from infection (Mongelli & Saleh, 2016). Mosquitoes employ an additional RNAi pathway which was previously associated primarily with TE silencing in Drosophila, as an antiviral defense system (Hess et al., 2011; Miesen, Joosten, & van Rij, 2016) (Kunitomi et al, submitted). The Piwi-interacting RNA (piRNA) pathway, mediated by Piwi proteins and associated 24-28nt small RNAs, involves the cleavage and processing of endogenous TE genomic antisense transcripts into small RNAs. These small RNAs target TE transcripts with appropriate sequence identity, yielding sense piRNAs that, in turn, drive further antisense transcript processing.

Transcripts derived from genomic EVE sequences can be processed into piRNAs, and a set of proteins responsible for their processing and maturation has been identified (Arensburger, Hice, Wright, Craig, & Atkinson, 2011; Goic, et al., 2016; Miesen, Girardi, & van Rij, 2015; Miesen, Ivens, Buck, & van Rij, 2016; Miesen, Joosten, et al., 2016; Varjak et al., 2017). Genomic EVE sequences confer resistance to viruses that encode identical sequences, in association with an accumulation of EVE-specific piRNAs (Kunitomi et al, submitted) (Miesen, Joosten, et al., 2016). Thus, the library of acquired viral sequences in the mosquito genome not only represents a record of the natural history of infection in this important vector species, but also a potential reservoir of immune memory. Understanding the acquisition of circulating viruses into these heritable genomic loci has major implications for mosquito immunity and disease transmission. Toward that end, recent publications have examined Flavivirus EVEs in both wild-caught mosquitoes and mosquito-derived cell lines (Suzuki et al., 2017) and nonretroviral EVEs across currently available genomic assemblies (Palatini et al., 2017). These studies have demonstrated that EVEs integrate in association with LTR sequences and integrate into genomic loci known as piClusters. However, because of difficulties resolving these genomic regions the full diversity of EVE sequences and their relationship to piRNAs derived from these sequences have not been yet described in a systematic way.

Here, we report the first such study, which examines the structure and genomic context of the collection of non-retroviral EVEs present in the *Aedes aegypti-derived* Aag2 cell line genome. Using an improved genomic assembly from long-read sequencing as the basis of our analysis, we characterize the structure and composition of EVE-containing loci across the entire genome. Additionally, we compare these data to small RNA sequencing data from Aag2 cells to assess the transcription and processing of small RNAs originating from these loci to understand the form and potential antiviral function of these important loci.

## RESULTS

### Sequencing and Assembly of the Aag2 Genome

Current assemblies of the *Ae. aegypti* genome are based on two sequencing strategies: one produced with the Illumina sequencing platform (hereto referred to as ‘UCB’) (Vicoso & Bachtrog, 2015) and one based on conventional Sanger sequencing (hereto referred to as ‘LVP’) (Nene et al., 2007). In both instances, the Liverpool strain of *Ae. aegypti* was sequenced (Table 1). A more recent study used Hi-C to further organize the Sanger-based *Ae. aegypti* assembly into chromosome level scaffolds (Dudchenko et al., 2017). Due to the highly repetitive nature of the *Ae. aegypti* genome and EVEs’ tendency to cluster with transposable elements within such repetitive regions, many EVEs are likely to be missing from the current *Ae. aegypti* assemblies, which are based on relatively short read lengths (Nene, et al., 2007; Vicoso & Bachtrog, 2015). Assessing the comprehensive genome-wide diversity and genomic context of EVE sequences therefore requires an improved genomic assembly. To this end, we employed single-molecule, real-time sequence technology (Pacific Biosystems) to generate a long read assembly of the genome of the cell line Aag2.

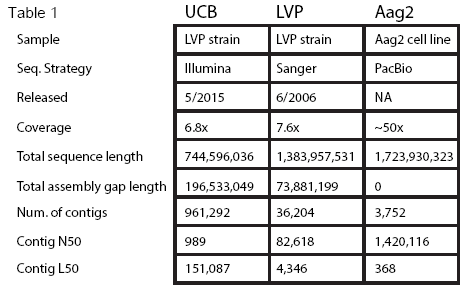

We achieved approximately 76-fold coverage of the *Ae. aegypti-based* Aag2 genome using the Single Molecule Real Time (SMRT) sequencing platform (P6/C4 chemistry) to shotgun sequence 116 SMRT cells generating 92.7 GB of sequencing data with an average read length of 15.5 kb. We used Falcon and Quiver to generate a *de novo* 1.7 Gbp assembly with a contig N50 of ~1.4 Mbp. Our draft assembly improves upon previous *Aedes* assemblies as measured by N50, L50, and by contig number (Table 1 and Figure 1a). A majority of the Aag2 assembly sequence is found on contigs 10-100x longer than previous assemblies. This increased contiguity allows the mapping of numerous contigs from the initial LVP assembly to single Aag2 contigs (SI Figure 1), and makes for an overall more ordered genome assembly.

**Figure 1.**
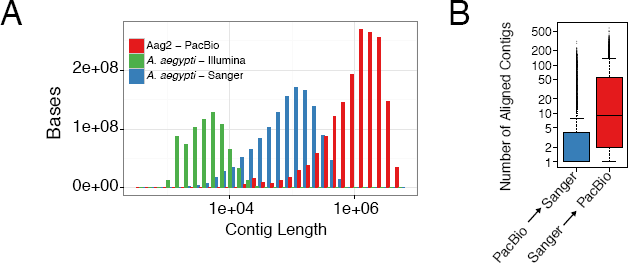
Contiguity of the *Aedes aegypti* genome is drastically improved in the Aag2 assembly. (A) Histogram of contig length vs. total amount of sequence contained in each bin. The Aag2 assembly achieved the largest contig sizes (by an order of magnitude) compared to previous *Aedes aegypti-derived* assemblies. This large contig size also resulted in more overall sequence information/number of bases. (B) Boxplots indicating number of contigs aligned between LVP (Sanger) and Aag2 (PacBio). When aligned to each other, more contigs from LVP are aligned to larger Aag2 contigs than vice versa.

### Repetitive nature of the Aag2 genome

The genome of *Ae. aegypti* was previously shown to contain high proportions of repeat-DNA (Nene, et al., 2007). The *Ae. aegypti-derived* Aag2 genome is no different, and is comprised of almost 55% repeat sequence (Table 1 and Table 2). Our sequencing strategy allows more repeats to be sequenced within a single read, and therefore better reflects the structure and organization of these repetitive elements. Direct alignment of contigs in the Aag2 assembly and those of previous *Ae. aegypti* assemblies reveal resolved rearrangements and distinct repeated regions that were collapsed into single sequences in the previous assemblies (Figure 1b and SI Figure 1). These regions can span 10-20kb (uncollapsed), illustrating the need for long read lengths to properly order the vast number of repetitive regions in the *Ae. aegypti* genome. Of these repetitive regions, over 75% are made up of transposon-derived sequence (Table 2).

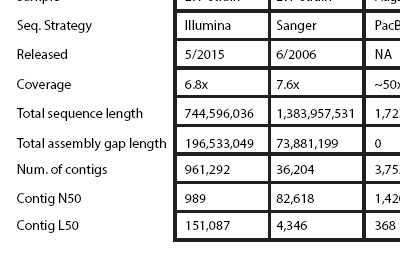

### Identification of EVEs in the Aag2 genome

Given their propensity to integrate into long, repetitive TE clusters (Figure 3c, 4b)(Parrish et al., 2015), our understanding of the EVE composition and structure has been limited. Our improved, long-read assembly can better define the complete set of EVEs contained within the Aag2 genome, hereby called the “EVEome”. Using a BLASTx-based approach, EVEs were identified with respect to each virus’ protein coding/(+)-sense strand (see Methods). We identified a total of 472 EVEs in our Aag2 genome assembly. These EVEs represented at least 8 annotated viral families, but were dominated by sequences derived from Rhabdoviridae, Flaviviridae, and Chuviridae. The identified EVEs covered 338,251 bp and ranged from 50 to 2,520 bp length with a median length of 620 bp (Figure 2a and b).

**Figure 2.**
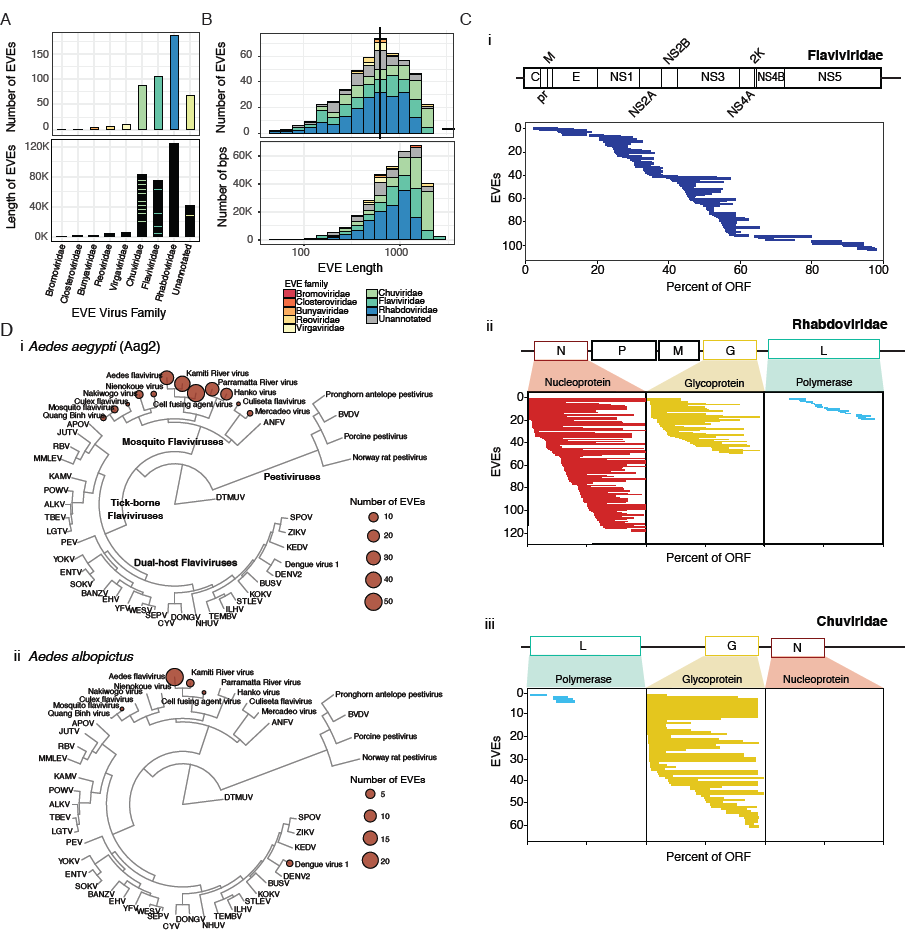
Identification of Endogenous Viral Elements (EVEs) in the Aag2 assembly. (A)Bar plots showing the number (top) and total length (bottom) of EVEs derived from each viral family. (B)Histogram showing the size distribution of EVEs in the Aag2 genome (top) and the total number of bases pairs derived from EVEs of a given size. The median EVE size (620bp) is indicated with a black bar. (C)Coverage plots of EVEs derived from the viral families (i) Flaviviridae, (ii) Rhabdoviridae, and (iii) Chuviridae. Each bar represents a single EVE, while its length and position denotes the region of the indicated ORF from which its sequence is derived. Length is expressed as a percentage of the total ORF to normalize for varying ORF lengths among different members of a given viral family. The genome organization of CFAV is presented for reference in (i). In (ii) and (iii), a generic genome is presented to illustrate from where EVEs are derived within the genome and within each specific ORF. (D)Phylogenetic relationship between 61 members of Flaviviridae. EVEs present in (i) *Ae. aegytpi* or (ii) *Ae. albopictus* which align to the indicated virus are marked with a colored circle. Size corresponds to abundance of EVEs derived from given species.

**Figure 3.**
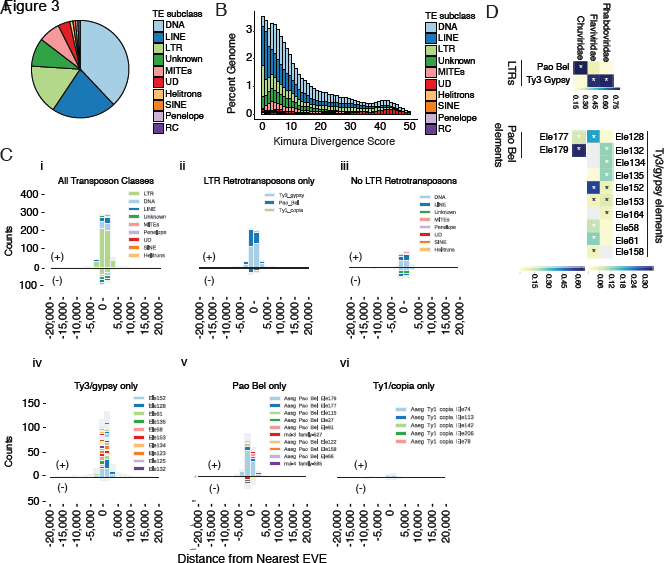
The repeat landscape of the *Aedes aegypti* genome is predominantly made up of transposable elements. (A)Pie chart representing relative numbers repetitive elements in the Aag2 genome. Further detail can be found in Table 2. (B)Stacked histogram of Kimura divergence for classes of TEs found in the Aag2 assembly, expressed as a function of percentage of the genome. A relatively recent expansion/active phase of LTRs is evident (increase in LTRs at low Kimura divergence scores). Kimura divergence scores are based on the accumulated mutations of a given TE sequence compared to a consensus. (C)Histograms showing counts of non-overlapping TEs closest to EVEs binned by distance, both upstream (negative x-axis values) and downstream (positive x-axis values). Positive y-axis counts refer to TE/EVE ‘pairs’ with the same strandedness, while negative y-axis counts are EVEs where the closest TE has the opposite strandedness. Total counts represented in each histogram: All classes (n=942); LTR only (n=614); No LTRs (n=328); Ty3/gypsy only (n=358); Pao Bel only (n=226); Ty1/copia only (n=30). (D)Heatmap showing categories of TEs nearest EVEs, categorized by the viral family from which the EVEs were derived. Only TEs with the same strandedness as its nearest EVE are shown. A “*” indicates significant enrichment by one-sided binomial test against the background prevalence of a given TE category in the genome (eg among all LTRs nearest Chuviridae-derived EVEs, Pao Bel elements are specifically enriched compared to the genome-wide counts of Pao Bel among all LTRs). Color indicates proportion of a given TE category nearest EVEs derived from the indicated viral family. Grey indicates the element was not found to be the closest TE to any EVEs derived from the indicated viral family. Only TE elements which made up at least 10% of the dataset for a given viral family are shown. “Pao Bel elements” refers to Chuviridae, while “Ty3/gypsy elements” corresponds to Flaviviridae and Rhabdoviridae. Total sample size of all TEs analyzed for each dataset: LTRs-Rhabdoviridae (n=130),Flaviviridae (n=181), Chuviridae (n=107); Pao Bel-Chuviridae (n=84); Ty3/gypsy-Rhabdoviridae (n=100), Flaviviridae (n=136).

To determine whether any region of the virus genome is more frequently acquired, EVEs were mapped onto the viral ORFs from which they derive. This analysis revealed asymmetric incorporation of certain viral ORFs (Figure 2c). *Flaviviridae-derived* EVEs (Figure 2c, i) primarily mapped toward the 5’ end of the single Flaviviral ORF, leaving a relative dearth of EVEs at the 3’ end. EVEs deriving from *Rhabdoviridae* primarily originate from the Nucleoprotein (N) and Glycoprotein (G) coding sequences, with only a few originating from the RNA-dependent RNA polymerase (L) (Figure 2c, i). The lack of EVEs mapping to the polymerase may be the result of RNA expression levels, with L being the least expressed gene (Conzelmann, 1998), suggesting that the template for cDNA synthesis is viral mRNA. The lack of EVEs originating from the Phosphoprotein (P) or Matrix protein (M) is more difficult to explain, potentially reflecting the availability of the RNA template for recombination, or deleterious effects associated with acquisition of these sequences. Interestingly, EVEs derived from Chuviridae primarily map to the ORF of the Glycoprotein. Given the diverse Chuviridae genome organization (which occur as unsegmented, bi-segmented, and possibly circularized negative-sense genomes)(Li et al., 2015), this pattern could also be the result of mRNA abundance, or other mechanistic peculiarities of Chuviridae interaction with EVE acquisition machinery (see below). Within each viral ORF, there is no obvious preference for EVEs from a specific location.

### Comparison of EVEs in *Ae. aegypti* and *Aedes albopictus* genomes

If EVEs serve as a representative record of viral infection over time, we hypothesized that EVEs present in two different species of mosquito would also differ (particularly given the relatively rare occurrence of genome fixation) (Holmes, 2011; Katzourakis & Gifford, 2010). The *Ae. aegypti* and *Ae. albopictus* species of mosquito occupy distinct (yet overlapping) regions around the globe (Kraemer, et al., 2015) and have, therefore, faced different viral challenges over time. While the EVEs present in the Aag2 and LVP *Ae. aegypti-based* genomes correspond well, *Ae. aegypti* and *Ae. albopictus* do not share any specific EVEs. However, exploring the Flaviviridae family of viruses in greater detail, the viral species from which these EVEs are derived do primarily overlap (Figure 2d). However, the relative abundance of EVEs derived from various viral species in *Ae. aegypti* and *Ae. albopictus* differs. The lack of specific EVEs in common between the mosquito genomes indicates EVE acquisition by *Ae. aegypti and Ae. albopictus* occurred post-speciation, an important factor when considering any differences in vector competence between these two species. However, it is also important to note that these species are estimated to have diverged around 71 million years ago (Chen et al., 2015), and so only under very strong and consistent positive selection could EVEs integrated pre-speciation have been preserved.

### Insights into the mechanism of EVE integration

Transposable elements (TEs) provide an important source of genomic variation that drives evolution by modifying gene regulation and genome organization, and through the acquisition of non-retroviral EVE sequences (Gifford, Pfaff, & Macfarlan, 2013; Thompson, Macfarlan, & Lorincz, 2016). Because of this relationship between transposable elements and EVE integration (Feschotte & Gilbert, 2012; Honda & Tomonaga, 2016; Miesen, Joosten, et al., 2016), and because our assembly is particularly suited for repeat identification and analysis, we explored the organization of TEs, specifically focusing on those proximal to EVE sequences.

The TEs in the Aag2 genome are derived from several different families (Fig 3a, S1 Fig). Kimura distribution analysis of TEs in the Aag2 genome can be used to ‘date’ the relative age of specific elements in the genome (Figure 3b) (Kimura, 1980). The distributions of Kimura scores for TEs in our assembly indicates relatively recent expansions of TEs, particularly LINE, LTR, and MITEs elements (Figure 3b; low Kimura scores indicate TEs that are closer to the element’s consensus sequence, while higher scores indicated more diverged TE sequences).

To determine whether a particular transposable element type is responsible for EVE integration, we identified TEs whose position directly overlaps EVE sequences (as called by RepeatMasker and BLASTX respectively). This approach identifies mobile elements most likely responsible for genomic integration of non-retroviral virus sequence. In line with observations in *Ae. albopictus (Chen, et al., 2015),* TEs overlapping EVE sequences are greatly enriched for LTR retrotransposons (S2 Fig (i); p-value < 3×10^^-60^). A similar pattern was observed when classifying the nearest upstream and downstream, non-overlapping TE sequences around each EVE (Figure 3c(i)). These results further implicate LTR retrotransposons in the acquisition of EVEs and indicate that the typical integration sites are composed of clusters of similar LTR retrotransposons.

Strikingly, 543 out of 614 LTRs shared the same polarity as their nearest-neighbor EVE (i.e. both TE and EVE elements are located on the same genomic DNA strand). These 543 LTRs made up the vast majority of all TEs with the same polarity as their nearest EVE (543/746; Figure 3c(i); p-value = 1.09x10^−252^ by one-sided binomial test). This bias is consistent with a copy-choice mechanism of recombination between LTR retrotransposon sequence and viral RNA (or viral mRNA) leading to EVE integration, as previously proposed (Cotton, Steinbiss, Yokoi, Tsai, & Kikuchi, 2016; Geuking et al., 2009). Our analysis of transposons in the Aag2 genome shows LTR-retrotransposons display less sequence diversity (by Kimura Divergence score; Figure 3b), indicating that they are currently (or were recently) actively replicating in the Aag2 cell line. Consistent with this idea, LTR-retrotransposon transcripts and proteins are readily detected in Aag2 cells (Maringer et al., 2017). These data are consistent with LTR-retrotransposons being responsible for the acquisition of the majority of EVEs observed in the *Ae. aegypti-derived* Aag2 genome.

Within the LTR retrotransposon family, both Ty3/gypsy and Pao Bel TEs are enriched surrounding EVEs (Figure 3c iv,v). Again, this enrichment for Ty3/gypsy and Pao Bel elements near EVE loci is strongest when the EVE and TE are in the same orientation (p-value = 6.90×10^−29^ and 1.71×10^−3^ respectively). The drastic bias in associated transposons based on directionality is not observed for other TE categories (Figure 3c, iii, vi). These data support Ty3/gypsy (and to a lesser extent Pao Bel) as the primary transposon type facilitating EVE genomic integration in the Aag2 cell line. Interestingly, an association between LTR Ty3/gypsy elements and integrated viral sequence has also been observed previously in plants (Lee, Nolan, Watson, & Tristem, 2013; Staginnus et al., 2007), suggesting a conserved mechanism for the acquisition of invading virus sequences and generation of EVEs.

EVE-proximal TEs of the Ty3/gypsy and Pao Bel families can be further partitioned into individual elements. Of these, many specific elements were enriched for being the nearest TE to an EVE (Figure 3c(iv, v)). Interestingly, EVEs derived from different virus families show different patterns of enrichment for nearby TEs (Figure 3d). Although Flaviviridae- and Rhabdoviridae-derived EVEs show strong enrichment for Ty3/gypsy transposable elements, Chuviridae-derived EVEs are associated with Pao Bel elements.

### EVEs associate with piClusters

The strong enrichment for multiple LTRs around EVEs (Fig 3c, S2 Fig) led us to examine the genomic context of EVE-TE integration sites in the genome. The large contig sizes associated with our long-read sequencing approach allow us to assess the large-scale spatial distribution of TEs and EVEs in the genome. Strikingly, we identified numerous loci where many EVE sequences overlapped with large regions of increased LTR density, some larger than 50kbp in length (Figure 4a). In some cases, these large loci are so densely packed with a single LTR they effectively “crowd out” any other repetitive elements (Figure 4b). Within these loci, EVE sequences are interspersed with TE fragments in unidirectional orientations (Figure 4c). They contain large numbers of EVEs derived from different viral families (Figure 4c-e) suggesting that these regions occasionally capture new TE-virus hybrids.

**Figure 4.**
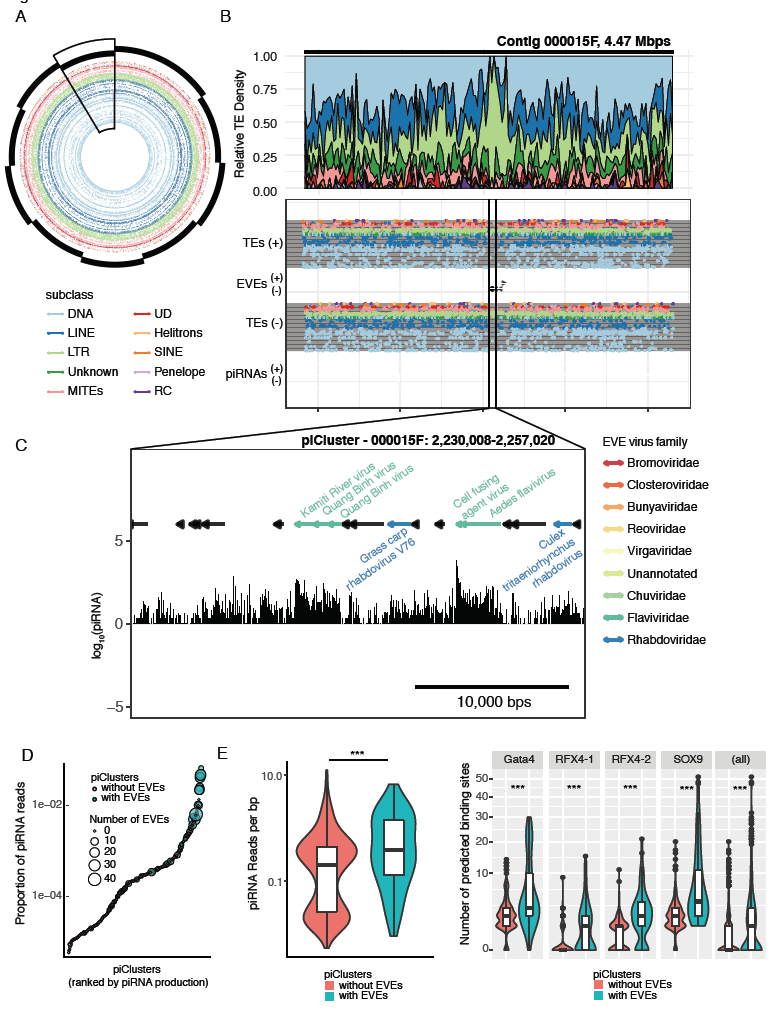
EVEs are primarily associated with LTR transposable elements. (A)Circle plot showing the arrangement and diversity of TE subclasses in the ten largest contigs in the Aag2 assembly. Individual contigs are denoted with staggered black bars. Specific TE elements are shown as dots, concentric rngs represent individual families within each TE subclass. Large scale fluctuations in TE density can be seen in specific contigs (contig 000015F, boxed). (B)Local density plots of a representative region of local LTR density in contig 000015F. (C)The regions of local LTR density corresponds to the location of numerous EVEs (black arrows) and piRNA density (bar chart, bottom track). Bioinformatically predicted piCluster corresponding to a portion of the large LTR density in contig 000015F. LTRs (shown as black bars) are interspersed with EVE sequences (colored by virus family). piRNA production (black bars, below) shows highest density in regions corresponding to EVE sequences. (D)Dotplot showing the relationship between piCluster EVE content and piRNA production. piClusters are ranked by piRNA production. (E)Violin plot comparing the distribution of piRNA density in piClusters with or without EVEs. (F)Violin plots comparing the number of predicted transcription factor binding sites in piClusters with or without EVEs.

The organization of these loci is similar to that of piClusters (Yamanaka, Siomi, & Siomi, 2014): piRNA-producing loci in the genome that result from the accumulation of TE fragments (due to non-random LTR integrase-directed integration)(Lesbats, Engelman, & Cherepanov, 2016). To assess the ability of these loci to produce piRNAs, we performed small RNA sequencing, employing a procedure to enrich for *bona fide* piRNAs. Indeed, we found that these loci produce a large number of piRNAs in a predominantly anti-sense orientation, consistent with the transcription of piClusters (Czech & Hannon, 2016; Yamanaka, et al., 2014).

We then used bioinformatic prediction to identify putative piClusters in the genome based upon piRNA mapping density. This analysis identified 469 piRNA-encoding loci (piClusters) using proTRAC (Rosenkranz, Rudloff, Bastuck, Ketting, & Zischler, 2015; Rosenkranz & Zischler, 2012), accounting for 5,774,304 bp (0.335%) of the genome. Depending on the mapping algorithm used, between 63% (bowtie) and 77% (sRNAmapper.pl, see Methods) of beta-eliminated small RNAs from Aag2 cells mapped to these loci. Of the identified piClusters, 65 (14.1%) have EVE sequences associated with them and 64 of these piCluster-resident EVE sequences act as the template for piRNAs. Of the 472 EVEs identified, 256 (66.7%) or 280,475 bp of the 411,239 EVE bp mapped to piClusters (68.2%, Fisher’s test p<2.2e-16, OR=203.42). Furthermore, a vast majority of piRNAs which map to EVEs are anti-sense to the coding sequence of the corresponding virus (544,429/547,014; 99.5%), meaning that majority of piRNAs produced from genomic EVEs are potentially antiviral.

To examine whether piCluster-resident EVEs are under selection we examined the relative transcription of piClusters throughout the genome. piClusters that contain EVEs tend to produce more piRNAs (Fig 4d,e) The increased piRNA production at EVE-containing loci, is consistent with the observation that piClusters with EVEs exhibit higher transcription factor binding site density (Fig 4f). GATA4, SOX9 and RFX transcription factor binding sites are all enriched near EVE-containing piClusters (p<2E-16, Wilcox rank-sum test). These data together suggest that selection acts at the level of EVE-specific piRNA production.

### piRNA abundance reflects the cellular immune state

EVEs, in combination with their associate piRNAs, make up a reservoir of small RNA immune memory. Notably, in the Aag2 cell line, piRNA production from EVEs derived from a given viral family does not completely correlate with genomic abundance of those EVEs, suggesting that the antiviral potential of the EVEome against a given viral family depends on the amount of viral genetic information stored in the host genome, the transcriptional activity of individual piClusters and the sequence identity of the resulting piRNAs to circulating viral challenges (Figure 5a,b).

**Figure 5.**
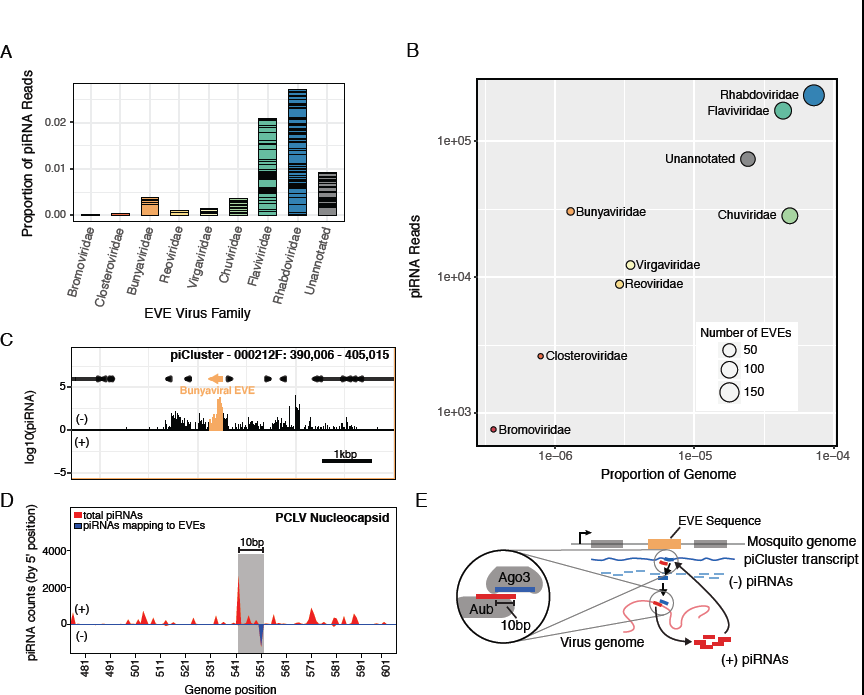
The antiviral potential of the cellular piRNA repertoire. (A)Bar plot showing the proportion of piRNA mapping to EVE sequences from a given viral family. Bars are split to show the relative contribution of specific EVE sequences. (B)Plot showing correlation the genomic footprint of EVEs from specific viral families in the Aag2 genome and their piRNA production. Bunyavirus and Chuvirus fall on opposite sides of the trend. (C)The EVE responsible for producing the anti-sense piRNA in (D). A spike of piRNAs is produced from the EVE (highlighted in orange) within the overall piCluster. (D)Mapping of cellular piRNAs to the bunyavirus PCLV genome reveals the pattern of piRNA processing. The sense-piRNA peak is offset by 10-bp from the antisense-piRNA peak (which also maps to an EVE within the Aag2 genome; blue line), showing a distinct ping-pong like pattern (highlighted by the grey rectangle. Interestingly, although the sequence of the antisense piRNAs map perfectly to the Aag2 genome/EVE, the sense piRNAs map perfectly to the PCLV virus sequence. piRNA counts are determined by the position of each piRNAs 5’ base. (E) Schematic showing the process of ping-pong piRNA amplification in the cell. An EVE sequence in a piCluster is transcribed to yield an antigenomic transcript. This transcript is processed into piRNAs which bind the genome of infecting viruses. Binding triggers processing of the viral genome into genomic piRNAs, which bind anti-genomic transcripts, leading to increased processing.

To examine the potential antiviral activity of piRNAs that originate from genomic EVEs, we mapped the same piRNA libraries (allowing for up to 3 mismatches) to contemporary viral genomes from which EVEs were expected to have derived (Figure 5d, S5 Fig). Aag2 cells are known to be persistently infected with cell-fusing agent virus (CFAV; *flaviviridae),* and were recently shown to also be persistently infected with Phasi Charoen-like virus (PCLV; *bunyaviridae)* (Maringer, et al., 2017) and these viruses, therefore, constitute potential substrates for recognition by EVE-derived piRNAs and subsequent processing. EVE-derived anti-genomic piRNAs (Figure 5c) only mapped to a single site on the PCLV nucleocapsid. However, we identified numerous sense piRNAs derived from PCLV, including a prominent peak which is offset from the EVE-derived anti-sense piRNA binding site by 10bps (Figure 5d,e). This pattern is consistent with EVE-derived piRNAs being funneled through canonical processing by the ping-pong amplification mechanism, being successfully loaded into the Piwi machinery and subsequently cleaving the viral mRNA. A similar pattern was observed for CFAV (Kunitomi, et al, submitted). Mapping of piRNAs to other viruses from which EVEs derived do not reveal this ‘response’ sense piRNA peak, presumably because those viruses are not currently replicating in the cells (S5 Fig). These observations indicate that an organism’s EVEome produces piRNAs capable of recognizing viruses and initiating an active response.

## DISCUSSION

A solid foundation with which to study the genetic factors contributing to vector competence is of utmost importance as arboviruses become an increasing burden globally. *In vivo* studies of mosquito immunity are a valuable but challenging approach to understanding arboviral life cycles. With this in mind, we generated a long-read assembly of the *Aedes aegypti* cell line, Aag2. With this highly contiguous assembly, we then identified nearly the entire set of endogenous viral elements and their surrounding genomic context in the Aag2 cell line at a genome-wide scale. The Aag2 cell line is an important model system for the characterization of arboviral replication in mosquito hosts. Considering the potential impact of EVE sequences and their associated piRNAs on viral infection, understanding the diversity of EVEs in commonly used cell lines is especially important.

Surveying the genome-wide collection of EVEs in the Aag2 genome provides not only a view of the historical interactions between host and virus, but also the repertoire of acquired sequences that define the piRNA-based immune system of this important model system. Our analysis refines our understanding of EVEs in the *Ae. aegypti* genome, their relationship to transposable elements, and the potential breadth of antiviral protection they provide. We propose that a mosquito’s EVEome, together with the piRNA system, represents a potentially long-lasting branch of its RNAi anti-viral defense system. Although all mosquito species share the same basic RNAi-based immune system, the differences in the EVEome of a given species, subpopulation, or individual, such as those observed between *Ae. aegypti* and *Ae. albopictus* (Figure 2a), may represent a factor contributing to inherent differences in vector competence across many different scales. Indeed, the EVEome of wild mosquito populations appears to be in rapid flux (Varjak, et al., 2017).

The presence of piRNA producing EVEs in the *Ae. aegypti* genome is reminiscent of the CRISPR system in bacteria. Both take advantage of the invading pathogen’s genetic material to create small RNAs capable of restricting an invading virus’ replication. Furthermore, both integrate into the host’s genome, potentially providing protection against infection across generations. Although the conservation of this pathway across Eukarya is not yet clear, recent publications have highlighted the integration of genetic material from non-retroviral RNA viruses into the genome of many different host species during infection. The antiviral activity of these sequences has not been established, however, a subset of EVEs found in mammalian systems seem to be under purifying selection, suggesting some potential benefit to the host (Horie, et al., 2010). In contrast to the evolutionary repurposing of retroviral sequences, the direct integration, transcription and processing of EVE sequences into antiviral small RNAs constitutes a mechanism by which these acquired sequences can be rapidly repurposed for host immune purposes.

Template switching during reverse transcription has previously been proposed to play a role in creating transposon-virus hybrids which integrate into the host genome to form EVEs (Cotton, et al., 2016; Geuking, et al., 2009). The apparent family-level specificity observed between Pao Bel TEs and Chuviridae-derived sequences and between Ty3/gypsy TEs and Flaviviridae and Rhabdoviridae sequences is interesting in this respect (Figure 3d). This could have occurred by chance, or may hint at an even deeper level of specificity directing capture of viral sequences by LTRs. Possibly these TEs and viruses share increased sequence homology leading to more frequent template switching (Delviks-Frankenberry et al., 2011), or replicate in a similar subcellular location. It is also possible that LTR/EVE pairs were selected for and maintained after EVE integration into the mosquito genome. Suzuki et al. note that in different strains of *Ae. albopictus,* the Flavivirus-derived EVEs are conserved, but their flanking regions can be quite distinct (Suzuki, et al., 2017). As the authors noted, this hints at an evolutionary role for the EVEs themselves, but not necessarily the specific surrounding regions. Given the difference in piRNA production among piClusters with EVEs and without (Figure 4d,e), selection may only act at the level of piRNA production, rather than the specific TE elements.

Uncovering the genomic context of EVEs highlights the potential for the piRNA system to shape the mosquito immune system. It also provides a foundation for future investigations into EVE function. Comparative genomic approaches that incorporate long-read sequencing to understand the diversity of the EVEome across populations will allow us to better understand the forces that underlie the epidemiology and population dynamics of arboviruses. Moreover, the potential to manipulate this heritable, anti-viral immune system could present opportunities for epidemiological interventions in natural settings, or as a genetic system to understand the insect immune system in the laboratory.

## Acknowledgements

We thank Dr. Kevin M. Dalton for helpful discussion and code for the analysis.

## MATERIALS AND METHODS

### Cells culture

*Aedes aegypti* Aag2 *(Lan & Fallon, 1990; Peleg, 1968)* cells were cultured at 28 °C without CO_2_ in Schneider’s *Drosophila* medium (GIBCO-Invitrogen), supplemented with 10% heat-inactivated fetal bovine serum (FBS), 1X non-essential amino acids (NEAA, UCSF Cell Culture Facility, 100X stock is 0.1 μM filtered, 10 mM each of Glycine, L-Alanine, L-Asparagine, L-Aspartic acid, L-Glutamic Acid, L-Proline, and L-Serine in de-ionized water), and 1X Penicillin-Streptomycin-Glutamine (Pen/Strep/G, 100X = 10,000 units of penicillin, 10,000 μg of streptomycin, and 29.2 mg/ml of L-glutamine, Gibco).

### DNA sequencing

Aag2 cells were grown in T-150 Flasks until ~80% confluent. Cells were then washed with dPBS twice and scrapped off in dPBS + 10 μg/ml RNase A (ThermoFisher). Genomic DNA (gDNA) was extracted from ~10ˆ8 Aag2 cells using the QIAamp DNA Mini Kit according to the manufacturer’s instructions with the optional RNase A treatment. Aag2 gDNA was re-suspended in 10mM Tris pH8, and the quality and quantity of the sample was assessed using the Agilent DNA12000 kit and 2100 Bioanalyzer system (Agilent Technologies), as well as the Qubit dsDNA Broad Range assay kit and Qubit Fluorometer (Thermo Fisher) and visualized by gel electrophoresis (1% TBE gel). After purification and quality control, a total of 130 ug of DNA was available for library preparation and sequencing.

SMRTbell libraries were prepared using Pacific Biosciences’ Template Prep Kit 1.0 (PacBio) and a slightly modified version of the Pacific Biosciences’ protocol, “Procedure & Checklist – 20-kb Template Preparation Using BluePippin Size-Selection System (15-kb Size Cutoff)”. Specifically, 52.5ug of gDNA were hydrodynamically sheared to target sizes of 30kb (26 μg) and 35 kb (26 μg) using the Megaruptor® (Diogenode) with long hydropores according to the manufacturer’s protocols. Size distributions of the final sheared gDNA were verified by pulse field electrophoresis of a 100ng sub-aliquot through 0.75% agarose using the Pippin Pulse (Sage Science), run according to the manufacturer’s “10-48 kb protocol” for 16 hrs. The two sheared samples were then pooled, for a total of 37ug sheared DNA to be used as input into SMRTbell preparation. Sheared DNA was subjected to DNA damage repair and ligated to SMRTbell adapters. Following ligation, extraneous DNA was digested with exonucleases and the resulting SMRTbell library was cleaned and concentrated with AMPure PB beads (Pacific Biosciences). A total of 20.5ug of library was available for size selection.

Approximately half (10ug) of the SMRTbell pooled SMRTbell library was size-selected using the BluePippin System (Sage Science) using a 15 kb cutoff and 0.75% agarose cassettes. To obtain longer read lengths, an additional 5ug of the library was selected using a 17kb cutoff.

Library quality and quantity were assessed using the Agilent 12000 DNA Kit and 2100 Bioanalyzer System (Agilent Technologies), as well as the Qubit dsDNA Broad Range Assay kit and Qubit Fluorometer (Thermo Fisher). An additional DNA Damage Repair step and AMPure bead cleanup were included after size-selection of the libraries.

Annealed libraries were then bound to DNA polymerases using 3nM of the SMRTbell library and 3X excess DNA polymerase at a concentration of 9nM using Pacific Biosciences DNA/Polymerase Binding Kit 1.0, (Pacific Biosciences). Bound libraries were sequenced on the Pacific Biosciences RSII using P6/C4 chemistry (PacBio), magnetic bead loading (PacBio) and 6 hour collection times. 84 SMRTcells of the > 15 kb library were loaded at concentrations of 75-100 pM on-plate. 32 SMRTcells of the > 17 kb library was prepared separately and loaded at on-plate concentrations of 40 pM and 60 pM. These 116 SMRTcells generated 92.7 GB of sequencing data, which resulted in approximately 76X coverage of the Aag2 genome. Average raw read length of 15.5KB, with average sub-reads length of 13.2kb. Assembly was performed using Quiver/FALCON

### Genome assembly statistics

Basic statistics (e.g. Size, Gaps, N50, L50, # contigs) for each genome analyzed was produced using Quast (Gurevich, Saveliev, Vyahhi, & Tesler, 2013).

As a complementary approach Benchmarking sets of Universal Single-Copy Orthologs (BUSCO) was also run using the Arthropod dataset in order to assess the completeness of genome assembly. Of the 2675 BUSCO groups searched only 81 were missing from the Aag2 assembly, indicating good assembly completeness. Of the 2315 BUSCOs found only 279 of them were annotated as fragmented, emphasizing the continuity of the assembly.

### Repeat Identification and Kimura Divergence

In order to *de novo* identify and classify novel repetitive elements from the Aag2 genome, RepeatModeler was run on the assembled genome using standard parameters. Outputs from RepeatModeler were cross-referenced with annotated entries for Aedes aegypti from TEfam. All entries from RepeatModeler that were >80% identical to TEfam entries were discarded as redundant. This combined annotated and de novo identified list of repeat elements was used to identify the genome wide occurrences of repeats using RepeatMasker using standard parameters.

Kimura scores and corresponding alignment information were extracted from the “.align” file as output by RepeatMasker. This information was used to make the stacked plot in figure 2 using R (version 3.30) and the ggplot2 package.

Citations:

Smit, AFA, Hubley, R. *RepeatModeler Open-1.0*.

2008-2015 <http://www.repeatmasker.org>

Smit, AFA, Hubley, R & Green, P. *RepeatMasker Open-4.0.* 2013-2015 <http://www.repeatmasker.org>.

Dr. Zhijian Jake Tu. TEfam. <http://tefam.biochem.vt.edu/tefam/index.php>

### EVE identification

Identification of EVEs was achieved using standalone Blast+ (Altschul, Gish, Miller, Myers, & Lipman, 1990). Blast Searches were run using the Blastx command specifying the genome as the query and a refseq library composed of the ssRNA and dsRNA viral protein-coding sequences from the NCBI genomes as the database. The E-value threshold was set at 10-6.

The EVE with the lower E-value was chosen for further analysis to predict EVEs that overlapped. Several Blast hits to viral protein genes were identified as artifacts because of their homology to eukaryotic genes (e.g. closteroviruses encode an Hsp70 homologue). These artifacts were filtered by hand.

### Identification of LTR enrichment near EVEs

Separate BED files containing all TEs in the Aag2 assembly and all EVEs in the Aag2 assembly were used as input to Bedtools *(bedtools closest* command using the *−io* flag, and *−id* or *−iu)* to find the single closest non-overlapping TE to each EVE (both upstream and downstream).

An in-house script compiled these two output files together and filtered them for the TE content of interest. TE categories (subclass, family, element) were assigned by RepeatMasker. Enrichment was compared to the prevalence of the TE element genome wide based on a one-sided binomial test. Stacked histograms were produced based on TE categories as found in Figure 3. The legend lists (up to) the 10 most prevalent TE elements of TE/EVE pairs in the same orientation. Plots were produced using Python (version 2.7.6) with the pandas and matplotlib plugins.

### Classification of nearest TE to EVEs by virus taxonomy

Taxonomy categories for viruses from which each EVEs derived were assigned using an in-house script. Assignments were made based on NCBI's taxonomy database (ftp://ftp.ncbi.nih.gov/pub/taxonomy/), with the following additional annotations by hand.

| Virus species | Assigned family |
| --- | --- |
| *Wuhan Mosquito Virus 8* | Chuviridae |
| *Wuchang Cockroach Virus 3* | Chuviridae |
| *Lishi Spider Virus 1* | Chuviridae |
| *Shayang Fly Virus 1* | Chuviridae |
| *Wenzhou Crab Virus 2* | Chuviridae |
| *Bole Tick Virus 2* | Rhabdoviridae |
| *Shayang Fly Virus 2* | Rhabdoviridae |
| *Wuhan Ant Virus* | Rhabdoviridae |
| *Wuhan Fly Virus 2* | Rhabdoviridae |
| *Wuhan House Fly Virus 1* | Rhabdoviridae |
| *Wuhan Mosquito Virus 9* | Rhabdoviridae |
| *Yongjia Tick Virus 2* | Rhabdoviridae |
| *Cilv-C* | Virgaviridae |
| *Citrus leprosis virus C* | Virgaviridae |
| *Blueberry necrotic ring blotch virus* | Virgaviridae |
| *Wutai Mosquito Virus* | Bunyaviridae |

Heat maps were produced using the Seaborn plugin for python. Only TEs with >=10% proportion in at least one sample (Flaviviridae, Chuviridae, or Rhabdoviridae) are shown. Color was assigned based on proportion of TE element/family in each viral category.

Enrichment was scored as above using a one-sided binomial test (significant is p-value < 0.0001).

### Small RNA bioinformatics

Adaptors were trimmed using Cutadapt (http://dx.doi.org/10.14806/ej.17.1.200) using the --discard-untrimmed and -m 19 flags to discard reads without adaptors and below 19 nt in length. Reads were mapped using bowtie (Langmead, Trapnell, Pop, & Salzberg, 2009)using the – v1 flag (− v3 in the case of Fig 5D and S5). Read distance overlaps were generated by viROME (Watson, Schnettler, & Kohl, 2013). Uniquely mapping piRNAs were used for Figure 4C (-m1 flag).

### piCluster Analysis

piClusters were identified using PROtrac (Rosenkranz & Zischler, 2012) based on mapping with positions for beta-eliminated small RNAs libraries from Aag2 cells from sRNAmapper.pl. Based on these predictions, visualizations of clusters were produced using EasyFig (Sullivan, Petty, & Beatson, 2011) for visualization of TEs and R for comparison of TEs, piRNA abundance and EVE positions.

### Sequence alignment and phylogenetic analysis

For phylogenetic analysis of Flaviviridae, polyprotein sequences from 61 members of the Flaviviridae family were aligned with MUSCLE (Edgar, 2004) and a maximum likelihood tree was generated with FastTree (Price, Dehal, & Arkin, 2009) using the generalised time reversible substitution model (“-gtr”). Trees were visualized and annotated with ggtree (DOI: 10.1111/2041-210X.12628).

### EVE coverage

Base R (version 3.3.0) was used to show regions individual EVEs span on the indicated viral family (and protein). EVE length is a function of the percentage of the respective ORF from which it derives.

### Data availability

The Aag2 genome (v 1.00) is available through VectorBase (https://www.vectorbase.org/organisms/aedes-aegypti/aag2/aag2).

Main datasets produced during this work have been provided in excel format.

---

**S1 Fig.**
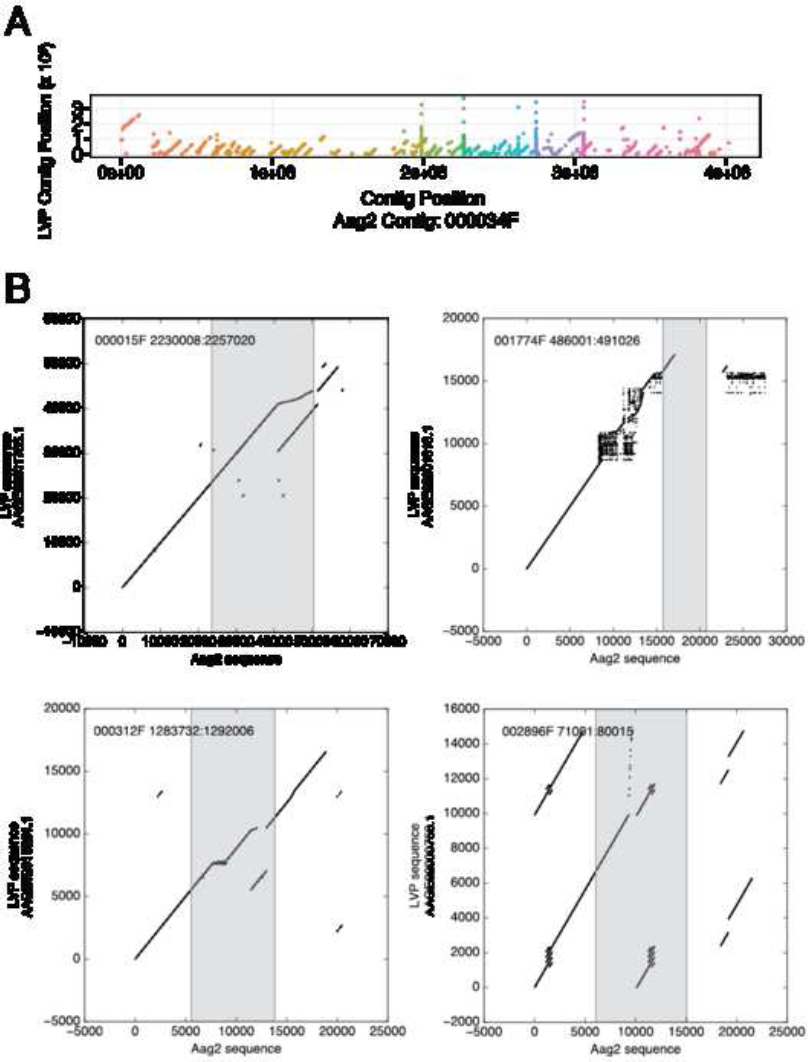
Transposons are distributed throughout the entire Aag2 genome. (A) Similar plot to Figure 2A, but showing all contigs of the Aag2 assembly. Circular plot of the Aag2 assembly, with every contig (black rectangles; ordered by size) and transposable element (circles; colored by TE class). Transposable elements are prevalent throughout the entire *Ae. aegypti* genome. Rectangles representing contigs are staggered to indicate relative contig size.

**S2 Fig.**
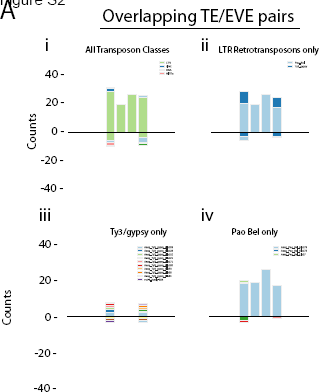
TEs which overlap EVEs are also overrepresented by LTR elements. (A) Histograms of TEs which overlap EVEs, broken down by the following categories. The left bin represents TEs whose start is upstream, and end overlaps the EVE. The 4th bin indicates TEs whose end is downstream, and start overlaps an EVE. The second bin indicates EVEs whose coordinates surround the TE. The third bin indicates TEs whose coordinates surround an EVE. Positive count values indicate TE and EVEs with shared directionality, while negative values represent TE and EVEs with opposite directionality. Some EVEs showed multiple overlapping TEs, all of which are represented on the charts.

**S3 Fig.**
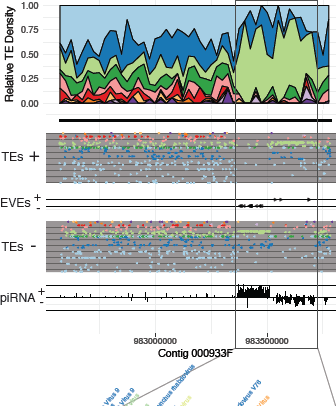
EVEs are typically found within unidirectional piRNA clusters. The left panels correspond to a region of Contig 000933F encoding 4 tandem, unidirectional piRNA clusters (as identified by proTRAC), each containing EVEs. Each cluster expresses piRNAs primarily anti-sense to the TEs/EVEs which define them. Similarly, a single large piRNA cluster on Contig 000044F is shown in the right panels. The shared directionality between TEs and EVEs (Figure 3B) is evident. Again, piRNA expression is almost exclusively in the antisense direction with respect to the TEs/EVEs.

**S4 Fig.**
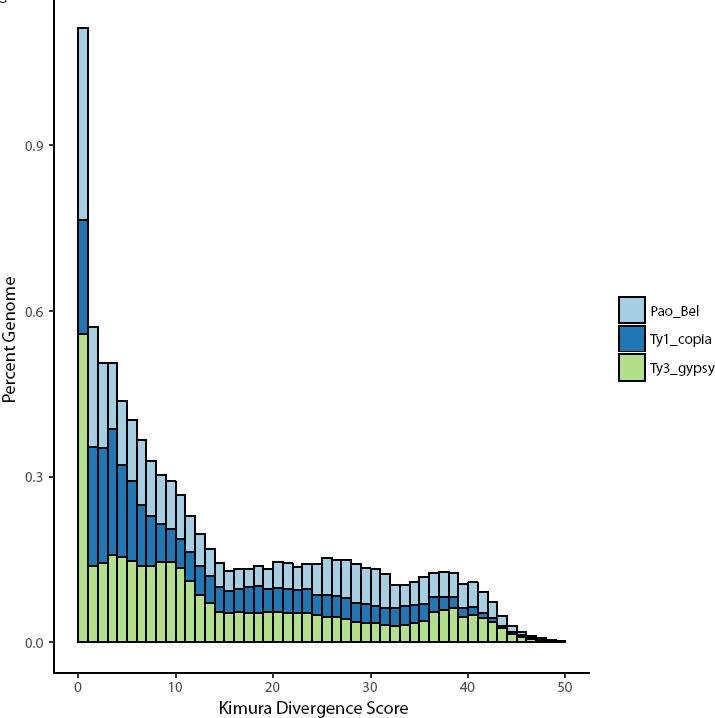
Kimura divergence scores of LTRs only show expansion of Pao Bel and Ty3/gypsy elements. Bar plot of kimura scores assigned to LTRs only, categorized by TE family and expressed as percent of total genome (as in Figure 2E). At very low (0-1) Kimura divergence scores, Ty3/gypsy and Pao Bel exhibit a marked increase in proportion of the genome.

**S5 Fig.**
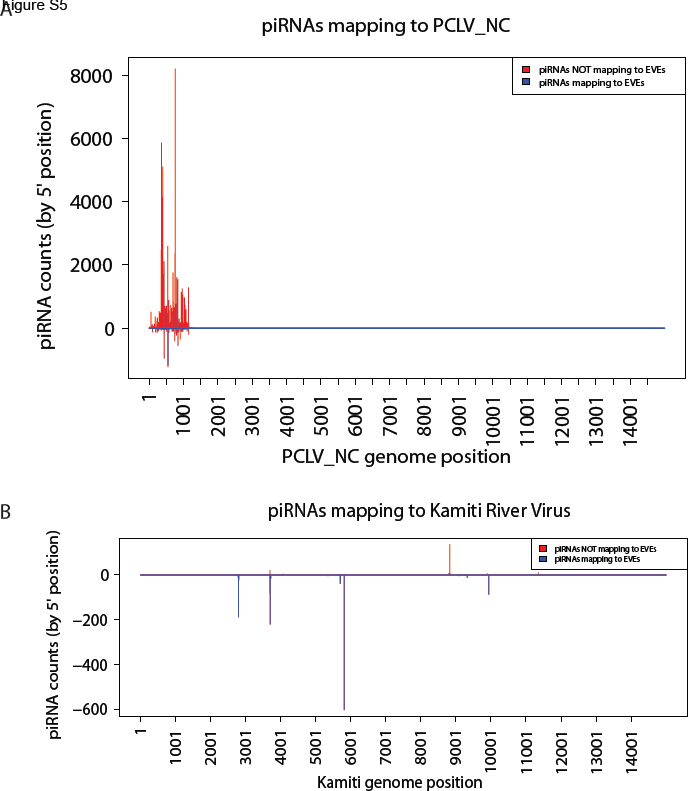
piRNA mapping to various viruses. (A) Same plot as Fig 5D, but showing the entire PCLV nucleocapsid region. (B) piRNAs mapping to Kamiti River Virus do not show the distinct ping-pong signature as seen for PCLV, despite a significant antisense piRNA peak deriving from an EVE.

## REFERENCES

Altschul, S. F., Gish, W., Miller, W., Myers, E. W., & Lipman, D. J. (1990). Basic local alignment search tool. J Mol Biol, 215(3), 403–410. doi:10.1016/S0022-2836(05)80360-2 S0022-2836(05)80360-2 [pii]

Arensburger, P., Hice, R. H., Wright, J. A., Craig, N. L., & Atkinson, P. W. (2011). The mosquito Aedes aegypti has a large genome size and high transposable element load but contains a low proportion of transposon-specific piRNAs. BMC Genomics, 12, 606. doi: 10.1186/1471-2164-12-606 1471 2164-12-606 [pii]

Aswad, A., & Katzourakis, A. (2012). Paleovirology and virally derived immunity. Trends Ecol Evol, 27(11), 627–636. doi: 10.1016/j.tree.2012.07.007 S0169-5347(12)00168-1 [pii]

Belyi, V. A., Levine, A. J., & Skalka, A. M. (2010a). Sequences from ancestral single-stranded DNA viruses in vertebrate genomes: the parvoviridae and circoviridae are more than 40 to 50 million years old. J Virol, 84(23), 12458–12462. doi: 10.1128/JVI.01789-10 JVI.01789-10 [pii]

Belyi, V. A., Levine, A. J., & Skalka, A. M. (2010b). Unexpected inheritance: multiple integrations of ancient bornavirus and ebolavirus/marburgvirus sequences in vertebrate genomes. PLoS Pathog, 6(7), e1001030. doi: 10.1371/journal.ppat. 1001030

Bennett, K. E., Olson, K. E., Munoz Mde, L., Fernandez-Salas, I., Farfan-Ale, J. A., Higgs, S., … Beaty, B. J. (2002). Variation in vector competence for dengue 2 virus among 24 collections of Aedes aegypti from Mexico and the United States. Am J Trop Med Hyg, 67(1), 85–92.

Bhatt, S., Gething, P. W., Brady, O. J., Messina, J. P., Farlow, A. W., Moyes, C. L., … Hay, S. I. (2013). The global distribution and burden of dengue. Nature, 496(7446), 504–507. doi: 10.1038/nature12060 nature12060 [pii]

Chen, X. G., Jiang, X., Gu, J., Xu, M., Wu, Y., Deng, Y., … James, A. A. (2015). Genome sequence of the Asian Tiger mosquito, Aedes albopictus, reveals insights into its biology, genetics, and evolution. Proc Natl Acad Sci USA, 112(44), E5907–5915. doi: 10.1073/pnas.1516410112 1516410112 [pii]

Chuong, E. B., Elde, N. C., & Feschotte, C. (2016). Regulatory evolution of innate immunity through co-option of endogenous retroviruses. Science, 351(6277), 1083–1087. doi: 10.1126/science.aad5497 351/6277/1083 [pii]

Conzelmann, K. K. (1998). Nonsegmented negative-strand RNA viruses: genetics and manipulation of viral genomes. Annu Rev Genet, 32, 123–162. doi: 10.1146/annurev.genet.32.1.123

Cotton, J. A., Steinbiss, S., Yokoi, T., Tsai, I. J., & Kikuchi, T. (2016). An expressed, endogenous Nodavirus-like element captured by a retrotransposon in the genome of the plant parasitic nematode Bursaphelenchus xylophilus. Sci Rep, 6, 39749. doi: 10.1038/srep39749 srep39749 [pii]

Czech, B., & Hannon, G. J. (2016). One Loop to Rule Them All: The Ping-Pong Cycle and piRNA-Guided Silencing. Trends Biochem Sci, 41(4), 324–337. doi: 10.1016/j.tibs.2015.12.008 S0968-0004( 15)00258-3 [pii]

Delviks-Frankenberry, K., Galli, A., Nikolaitchik, O., Mens, H., Pathak, V. K., & Hu, W. S. (2011). Mechanisms and factors that influence high frequency retroviral recombination. Viruses, 3(9), 1650–1680. doi: 10.3390/v3091650 viruses-03-01650 [pii]

Dudchenko, O., Batra, S. S., Omer, A. D., Nyquist, S. K., Hoeger, M., Durand, N. C., … Aiden, E. L. (2017). De novo assembly of the Aedes aegypti genome using Hi-C yields chromosome-length scaffolds. Science. doi: eaal3327 [pii] 10.1126/science.aal3327 science.aal3327 [pii]

Edgar, R. C. (2004). MUSCLE: multiple sequence alignment with high accuracy and high throughput. Nucleic Acids Res, 32(5), 1792–1797. doi: 10.1093/nar/gkh340 32/5/1792 [pii]

Feschotte, C., & Gilbert, C. (2012). Endogenous viruses: insights into viral evolution and impact on host biology. Nat Rev Genet, 13(4), 283–296. doi: 10.1038/nrg3199 nrg3199 [pii]

Geuking, M. B., Weber, J., Dewannieux, M., Gorelik, E., Heidmann, T., Hengartner, H., … Hangartner, L. (2009). Recombination of retrotransposon and exogenous RNA virus results in nonretroviral cDNA integration. Science, 323(5912), 393–396. doi: 10.1126/science.1167375 323/5912/393 [pii]

Gifford, W. D., Pfaff, S. L., & Macfarlan, T. S. (2013). Transposable elements as genetic regulatory substrates in early development. Trends Cell Biol, 23(5), 218–226. doi: 10.1016/j.tcb.2013.01.001 S0962-8924( 13)00003-2 [pii]

Gilbert, C., & Feschotte, C. (2010). Genomic fossils calibrate the long-term evolution of hepadnaviruses. PLoS Biol, 8(9). doi: 10.1371/journal.pbio.1000495 e1000495 [pii]

Goic, B., Stapleford, K. A., Frangeul, L., Doucet, A. J., Gausson, V., Blanc, H., … Saleh, M. C. (2016). Virus-derived DNA drives mosquito vector tolerance to arboviral infection. Nat Commun, 7, 12410. doi: 10.1038/ncomms12410 ncomms12410 [pii]

Goic, B., Vodovar, N., Mondotte, J. A., Monot, C., Frangeul, L., Blanc, H., … Saleh, M. C. (2013). RNA-mediated interference and reverse transcription control the persistence of RNA viruses in the insect model Drosophila. Nat Immunol, 14(4), 396–403. doi: 10.1038/ni.2542 ni.2542 [pii]

Gurevich, A., Saveliev, V., Vyahhi, N., & Tesler, G. (2013). QUAST: quality assessment tool for genome assemblies. Bioinformatics, 29(8), 1072–1075. doi: 10.1093/bioinformatics/btt086 btt086 [pii]

Hess, A. M., Prasad, A. N., Ptitsyn, A., Ebel, G. D., Olson, K. E., Barbacioru, C., … Campbell, C. L. (2011). Small RNA profiling of Dengue virus-mosquito interactions implicates the PIWI RNA pathway in anti-viral defense. BMC Microbiol, 11, 45. doi: 10.1186/1471-2180-11-45 1471 2180-11-45 [pii]

Holmes, E. C. (2011). The evolution of endogenous viral elements. Cell Host Microbe, 10(4), 368–377. doi: 10.1016/j.chom.2011.09.002 S1931-3128(11)00285-X [pii]

Honda, T., & Tomonaga, K. (2016). Endogenous non-retroviral RNA virus elements evidence a novel type of antiviral immunity. Mob Genet Elements, 6(3), e1165785. doi: 10.1080/2159256X.2016.1165785 1165785 [pii]

Horie, M., Honda, T., Suzuki, Y., Kobayashi, Y., Daito, T., Oshida, T., … Tomonaga, K. (2010). Endogenous non-retroviral RNA virus elements in mammalian genomes. Nature, 463(7277), 84–87. doi: 10.1038/nature08695 nature08695 [pii]

Katzourakis, A., & Gifford, R. J. (2010). Endogenous viral elements in animal genomes. PLoS Genet, 6(11), e1001191. doi: 10.1371/journal.pgen.1001191

Kimura, M. (1980). A simple method for estimating evolutionary rates of base substitutions through comparative studies of nucleotide sequences. J Mol Evol, 16(2), 111–120.

Kraemer, M. U., Sinka, M. E., Duda, K. A., Mylne, A. Q., Shearer, F. M., Barker, C. M., … Hay, S. I. (2015). The global distribution of the arbovirus vectors Aedes aegypti and Ae. albopictus. Elife, 4, e08347. doi: 10.7554/eLife.08347

Kramer, L. D. (2016). Complexity of virus-vector interactions. Curr Opin Virol, 21, 81–86. doi: S1879-6257(16)30104-3 [pii] 10.1016/j.coviro.2016.08.008

Kramer, L. D., & Ciota, A. T. (2015). Dissecting vectorial capacity for mosquito-borne viruses. Curr Opin Virol, 15, 112–118. doi: 10.1016/j.coviro.2015.10.003 S1879-6257(15)00153-4 [pii]

Lan, Q., & Fallon, A. M. (1990). Small heat shock proteins distinguish between two mosquito species and confirm identity of their cell lines. Am J Trop Med Hyg, 43(6), 669–676.

Langmead, B., Trapnell, C., Pop, M., & Salzberg, S. L. (2009). Ultrafast and memory- efficient alignment of short DNA sequences to the human genome. Genome Biol, 10(3), R25. doi: 10.1186/gb-2009-10-3-r25 gb-2009-10-3-r25 [pii]

Lee, A., Nolan, A., Watson, J., & Tristem, M. (2013). Identification of an ancient endogenous retrovirus, predating the divergence of the placental mammals. Philos Trans R Soc Lond B Biol Sci, 368(1626), 20120503. doi: 10.1098/rstb.2012.0503 rstb.2012.0503 [pii]

Lesbats, P., Engelman, A. N., & Cherepanov, P. (2016). Retroviral DNA Integration. Chem Rev, 116(20), 12730–12757. doi: 10.1021/acs.chemrev.6b00125

Li, C. X., Shi, M., Tian, J. H., Lin, X. D., Kang, Y. J., Chen, L. J… Zhang, Y. Z. (2015). Unprecedented genomic diversity of RNA viruses in arthropods reveals the ancestry of negative-sense RNA viruses. Elife, 4.doi: 10.7554/eLife.05378

Maringer, K., Yousuf, A., Heesom, K. J., Fan, J., Lee, D., Fernandez-Sesma, A., … Davidson, A. D. (2017). Proteomics informed by transcriptomics for characterising active transposable elements and genome annotation in Aedes aegypti. BMC Genomics, 18(1), 101. doi: 10.1186/s12864-016-3432-5 10.1186/s12864-016-3432-5 [pii]

Miesen, P., Girardi, E., & van Rij, R. P. (2015). Distinct sets of PIWI proteins produce arbovirus and transposon-derived piRNAs in Aedes aegypti mosquito cells. Nucleic Acids Res, 43(13), 6545–6556. doi: 10.1093/nar/gkv590 gkv590 [pii]

Miesen, P., Ivens, A., Buck, A. H., & van Rij, R. P. (2016). Small RNA Profiling in Dengue Virus 2-Infected Aedes Mosquito Cells Reveals Viral piRNAs and Novel Host miRNAs. PLoS Negl Trop Dis, 10(2), e0004452. doi: 10.1371/journal.pntd.0004452 PNTD-D-15-01748 [pii]

Miesen, P., Joosten, J., & van Rij, R. P. (2016). PIWIs Go Viral: Arbovirus-Derived piRNAs in Vector Mosquitoes. PLoS Pathog, 12(12), e1006017. doi: 10.1371/journal.ppat. 1006017 PPATHOGENS-D-16-02114 [pii]

Mongelli, V., & Saleh, M. C. (2016). Bugs Are Not to Be Silenced: Small RNA Pathways and Antiviral Responses in Insects. Annu Rev Virol, 3(1), 573–589. doi: 10.1146/annurev-virology-110615-042447

Nene, V., Wortman, J. R., Lawson, D., Haas, B., Kodira, C., Tu, Z. J., … Severson, D. W. (2007). Genome sequence of Aedes aegypti, a major arbovirus vector. Science, 316(5832), 1718–1723. doi: 1138878 [pii] 10.1126/science. 1138878

Palatini, U., Miesen, P., Carballar-Lejarazu, R., Ometto, L., Rizzo, E., Tu, Z., … Bonizzoni, M. (2017). Comparative genomics shows that viral integrations are abundant and express piRNAs in the arboviral vectors Aedes aegypti and Aedes albopictus. BMC Genomics, 18(1), 512. doi: 10.1186/s12864-017-3903-3 10.1186/s12864-017-3903-3 [pii]

Parrish, N. F., Fujino, K., Shiromoto, Y., Iwasaki, Y. W., Ha, H., Xing, J., … Tomonaga, K. (2015). piRNAs derived from ancient viral processed pseudogenes as transgenerational sequence-specific immune memory in mammals. RNA, 21(10), 1691–1703. doi: 10.1261/rna.052092.115 rna.052092.115 [pii]

Peleg, J. (1968). Growth of arboviruses in monolayers from subcultured mosquito embryo cells. Virology, 35(4), 617–619.

Price, M. N., Dehal, P. S., & Arkin, A. P. (2009). FastTree: computing large minimum evolution trees with profiles instead of a distance matrix. Mol Biol Evol, 26(7), 1641–1650. doi: 10.1093/molbev/msp077 msp077 [pii]

Rosenkranz, D., Rudloff, S., Bastuck, K., Ketting, R. F., & Zischler, H. (2015). Tupaia small RNAs provide insights into function and evolution of RNAi-based transposon defense in mammals. RNA, 21(5), 911–922. doi: 10.1261/rna.048603.114 rna.048603.114 [pii]

Rosenkranz, D., & Zischler, H. (2012). proTRAC‐‐a software for probabilistic piRNA cluster detection, visualization and analysis. BMC Bioinformatics, 13, 5. doi: 10.1186/1471-2105-13-5 1471-2105-13-5 [pii]

Staginnus, C., Gregor, W., Mette, M. F., Teo, C. H., Borroto-Fernandez, E. G., Machado, M. L., … Schwarzacher, T. (2007). Endogenous pararetroviral sequences in tomato (Solanum lycopersicum) and related species. BMC Plant Biol, 7, 24. doi: -1471-2229‐-7-24 [pii] 10.1186/1471-2229-7-24

Sullivan, M. J., Petty, N. K., & Beatson, S. A. (2011). Easyfig: a genome comparison visualizer. Bioinformatics, 27(7), 1009–1010. doi: 10.1093/bioinformatics/btr039 btr039 [pii]

Suzuki, Y., Frangeul, L., Dickson, L. B., Blanc, H., Verdier, Y., Vinh, J., … Saleh, M. C. (2017). Uncovering the repertoire of endogenous flaviviral elements in Aedes mosquito genomes. J Virol, doi: JVI.00571-17 [pii] 10.1128/JVI.00571-17

Taylor, D. J., Leach, R. W., & Bruenn, J. (2010). Filoviruses are ancient and integrated into mammalian genomes. BMC Evol Biol, 10, 193. doi: 10.1186/1471-2148-10¬193 1471-2148-10-193 [pii]

Thompson, P. J., Macfarlan, T. S., & Lorincz, M. C. (2016). Long Terminal Repeats: From Parasitic Elements to Building Blocks of the Transcriptional Regulatory Repertoire. Mol Cell, 62(5), 766–776. doi: 10.1016/j.molcel.2016.03.029 S1097-2765(16)30012-0 [pii]

Varjak, M., Maringer, K., Watson, M., Sreenu, V. B., Fredericks, A. C., Pondeville, E., … Schnettler, E. (2017). Aedes aegypti Piwi4 Is a Noncanonical PIWI Protein Involved in Antiviral Responses. mSphere, 2(3). doi: e00144-17 [pii] 10.1128/mSphere.00144-17 mSphere00144-17 [pii]

Vicoso, B., & Bachtrog, D. (2015). Numerous transitions of sex chromosomes in Diptera. PLoS Biol, 13(4), e1002078. doi: 10.1371/journal.pbio.1002078 PBIOLOGY-D-14-02861 [pii]

Watson, M., Schnettler, E., & Kohl, A. (2013). viRome: an R package for the visualization and analysis of viral small RNA sequence datasets. Bioinformatics, 29(15), 1902–1903. doi: 10.1093/bioinformatics/btt297 btt297 [pii]

Yamanaka, S., Siomi, M. C., & Siomi, H. (2014). piRNA clusters and open chromatin structure. Mob DNA, 5, 22. doi: 10.1186/1759-8753-5-22 1759-8753-5-22 [pii]

